# Efficient Endogenous Tagging in the Sea Urchin, *Lytechinus pictus*, Using CRISPR/Cas9-mediated Split-Fluorescent Protein Knock-In

**DOI:** 10.64898/2026.07.06.736833

**Authors:** Yoon Lee, Chloe Jenniches, Svenja Kling, Evan Tjeerdema, Elliot Jackson, Alexandre Paix, Amro Hamdoun

## Abstract

Precise knock-in of fluorescent reporters is a powerful tool for studying the dynamic cellular and molecular processes of embryogenesis. However, conventional CRISPR-Cas9 knock-in of large inserts, such as full-length fluorescent proteins, is inefficient. This has limited its application in many emerging model systems, including sea urchins. Here, we overcome this barrier using a transgenic *Lytechinus pictus* line that constitutively and ubiquitously expresses a large fragment of mNeonGreen (mNG3K^1-10^). In this line, fluorescence is only reconstituted when CRISPR-mediated knock-in delivers mNG2^11^, the 11th beta strand of the fluorescent protein, to complement the constitutively expressed fragment. Because this strategy requires integrating only the short 11th-strand, together with short homology arms (∼130 nt total), by homology directed repair, it circumvents the size constraints that limit conventional full-length reporter knock-ins using CRISPR. Using this approach, we achieved integration efficiencies of 14-22%, roughly an order of magnitude higher than those obtained with full-length fluorescent protein knock-ins. This provides a streamlined, scalable method for endogenous protein visualization in echinoderm embryos and a valuable resource for studying gene function, morphogenesis, and toxicant response in this classic developmental model.

## Introduction

Transgenic technologies have been instrumental to unlocking the biology of a diverse range of emerging animal models. The most commonly used technologies broadly fall into two classes, namely being the use of either integrases or nucleases. The first typically drives semi-random integrations of the desired cargo into the genome. However, it does not directly label target proteins, and it depends upon exogenous promoters. The second approach is the use of programmable nucleases, which resolve these limitations by enabling site-specific modifications at genomic loci. CRISPR-Cas9 is the most widely used of these (Boel et al., 2018; Lin et al., 2014; Paix et al., 2015; Seleit et al., 2021), because of its target flexibility, low cost and specificity (Jinek et al., 2012). However, despite its potential, CRISPR-Cas9 knock-in of fluorescent reporters in emerging models remains underutilized due to its low efficiency.

Sea urchins are a classical developmental that have recently crossed the barrier from transient to stable genetics (Jackson et al., 2024; Vyas et al., 2022). The first adaptation of CRISPR knock-in (Oulhen et al., 2023) in sea urchins was performed in *Strongylocentrotus purpuratus*, targeting the polyketide synthetase gene (PKS1). PKS1 is an enzyme involved in the production of echinochrome within pigment cells (Castoe et al., 2007) during early larval development, and has been used for proof-of-principle studies to test gene-editing technology, due to the easily observable albinism produced when it is interrupted (Liu et al., 2019; Yaguchi et al., 2020). In this prior study, a double-stranded (ds) biotinylated donor template with short homology arms (30 - 40 bp) containing the full length coding sequence for the fluorescent protein mNeonGreen was inserted into the endogenous coding region, resulting in mosaic somatic reporter expression in approximately 2% of injected F0 embryos. Since germline integration would only occur in a fraction of the animals produced by this method, this level of efficiency is impractical for generation of stable transgenic lines.

A major factor contributing to the inefficiency of CRISPR-Cas9-mediated knock-in is the size of the insert in the donor template sequence. Several modifications/optimizations have been proposed to address this issue, including the biotinylation of donor templates, long homology arms, and use of the single stranded donors (Paix et al., 2017). Despite these modifications, cargo size remains a fundamental limitation. Integration of large DNA cargos, such as full-length fluorescent protein, is less efficient than the insertion of small epitope tags (Paix et al., 2017).

Split fluorescent proteins (split-FPs) have emerged as a transformative strategy to overcome the limitation of insert size. Split-FPs are divided into two fragments that reconstitute a functional fluorescent protein when brought into close proximity (Cabantous et al., 2005). Most commonly, the two fragments consist of the first ten domains of the fluorescent protein (FP^1-10^) and the eleventh domain (FP^11^) (Cabantous et al., 2005; Feng et al., 2017; Tamura et al., 2021; Zhou et al., 2020). Less commonly used are pairs of other domains (Gaglia and Lahav, 2014; Shyu et al., 2006). This technology has been widely leveraged to improve the efficiency of generating transgenic lines in animal models by using CRISPR-Cas9 to insert the sequence encoding the smaller fluorescent protein fragment (∼48-60 nt) at the N- or C-terminus of an endogenous gene, while using other methods (such as transposase-mediated genomic insertion) to integrate the complementary larger fluorescent protein fragment required for fluorescence reconstitution (Goudeau et al., 2021; Kamiyama et al., 2021; Ligunas et al., 2024; O’Hagan et al., 2021).

Because only a short DNA sequence is inserted at the target locus, donor template design is simplified and homology-directed repair efficiency is increased compared to full-length fluorescent protein knock-ins. In organisms such as *Caenorhabditis elegans* (worm) (Goudeau et al., 2021), *Drosophila melanogaster* (fly) (Kamiyama et al., 2021), *Mus musculus* (mouse) (O’Hagan et al., 2021), *Danio rerio* (fish) (Ligunas et al., 2024) knock-in of short tags such as mNG2^11^, GFP^11^, and other variants using single stranded oligonucleotide donors (ssODN) has yielded significantly higher rates of successful integration compared to full-length tags. This enables sensitive detection of protein expression only when the tagged protein is produced.

Recently, a transposon-based approach was successfully adapted for use in echinoderms enabling genome integration in sea urchins (Jackson et al., 2024). Here, we build upon this to generate a *Lytechinus pictus* transgenic line expressing a transgene encoding the sequence for a plasma membrane localizing (LCK) Electra2 (blue fluorescent protein; (Papadaki et al., 2022)) and mNG3K^1-10^ (first ten domains of mNeonGreen; (Zhou et al., 2020)) separated by a tandem P2A and T2A (PT2A) peptide cleavage sequence; which we term Electra Neon Green 3K (ENG3K). In this study we generated sea urchins in which ENG3K was driven by the *Lytechinus variegatus* polyubiquitin-C (Entrez ID: LOC121415894) promoter, capable of driving the constitutive and ubiquitous expression of the large mNG3K^1-10^ across many spatial domains. The resulting line is referred to as Lv-polyubiquitin-C::LCK-Electra2_PT2A_mNG3K^1-10^ (abbreviated as LvP::ENG3K).

To evaluate the performance and generalizability of this system, we targeted multiple endogenous genes representing distinct cell types and compared CRISPR-Cas9 knock-in efficiencies across loci. We demonstrate that this founder line enables visualization of endogenous proteins following CRISPR-mediated insertion of the complementary mNG2^11^ sequence into target genes using small donor templates.

## Results

To determine the optimal strategy for improving CRISPR-Cas9 knock-in efficiency in sea urchin embryos, we compared several commonly proposed donor template optimizations. These included the biotinylation of donor templates, short homology arms, and the use of single-stranded donors. We measured knock-in efficiency across these four different donor templates, targeting the endogenous *FoxA* locus (Figure 1), using transient (mRNA overexpression) methods. The four donor templates included dsDNA mNeonGreen (714 nt), biotinylated dsDNA mNeonGreen (714 nt), ssDNA mNeonGreen (714 nt), and split mNG donor templates (54 nt). While unmodified dsDNA, biotinylated dsDNA, and ssDNA donors encoding full length mNeonGreen produced low knock-in efficiencies (2.2%, 0.4%, and 2.4% respectively). In contrast, the split-FP donor template yielded a significant increase in knock-in efficiency, reaching approximately 15-25% of injected embryos (p < 0.05 across all comparisons). Data for each replicate performed for these comparisons are available in Supplementary Table S1. These results supported the notion that donor template size was the most significant factor impacting knock-in efficiency.

**Figure 1.**
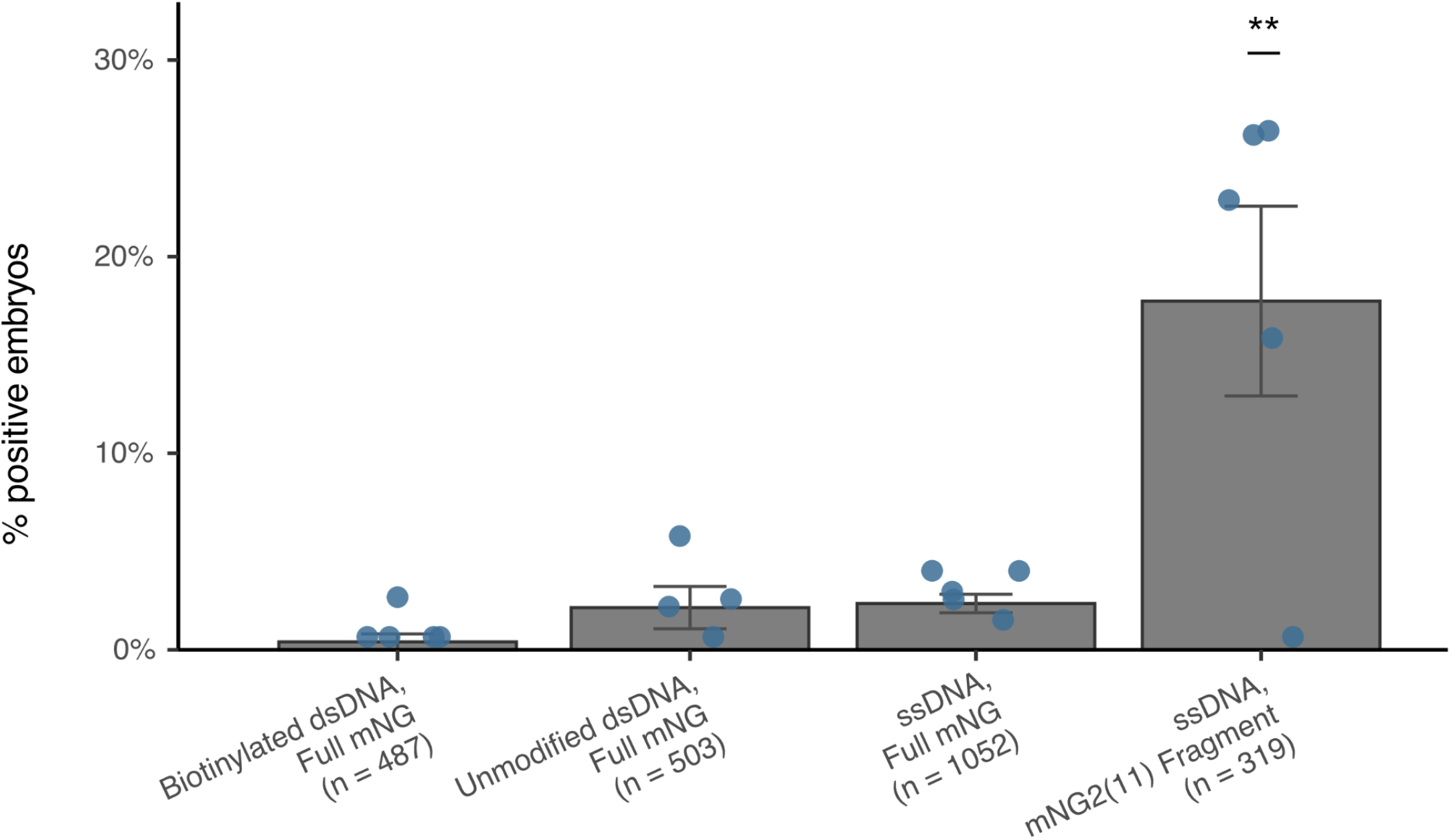
Short ssDNA donor templates significantly improve CRISPR-Cas9 knock-in efficiency. The efficiency of four different CRISPR-Cas9 HDR mediated knock-in strategies were compared by targeting the transcription factor FoxA. Embryos were injected with one of four donor types: biotinylated double-stranded DNA (dsDNA) encoding full-length mNeonGreen (mNG), unmodified dsDNA encoding full-length mNG, single-stranded DNA (ssDNA) encoding full-length mNG, or ssDNA encoding the split mNG2^11^ fragment. Bars show the mean percentage of positive embryos per donor type; error bars denote the standard error of the mean (SEM); overlaid points are individual biological replicates (n = 5 per condition; total embryos scored = 487, 503, 1,052, and 319, respectively, in the order shown). Conditions were compared by one-way ANOVA with Tukey’s HSD post-hoc test. The ssDNA mNG2^11^-fragment donor yielded a significantly higher proportion of positive embryos than the biotinylated dsDNA (p = 0.0012), unmodified dsDNA (p = 0.0049), and ssDNA full-length mNG (p = 0.0034) donors; no other pairwise comparison was significant (** p < 0.01).

In order to utilize the split-FP approach in a stable genetic line, the mNG3K^1-10^ fragment needs to be either expressed constitutively or during the same developmental time frame for a protein of interest that is tagged with mNG2^11^. We adopted the first approach to provide the most flexibility and utility across many gene targets. Since a transgenic line only expressing mNG3K^1-10^ is not possible to visually screen for integration (since mNG3K^1-10^ alone is not fluorescent) we generated a bicistronic construct, pUC19-LvP::ENG3K, containing both mNG3K^10^ (Ex λ 506 nm/Em λ 517 nm; (Shaner et al., 2013; Zhou et al., 2020)) and a spectrally distinct reporter LCK-Electra2 (Ex λ 403 nm/Em 454 λ; (Papadaki et al., 2022)) allowing for visual screening.

Three important considerations were made in our construct design (Figure 2A). First, within the sequence of the ENG3K transgene, the sequence for LCK-Electra2 is separated from the sequence for mNG3K^1-10^ by a PT2A peptide cleavage sequence to ensure that translated mNG3K^1-10^ is separated from the plasma membrane localizing LCK-Electra2 and free to complement mNG2^11^-tagged proteins across subcellular domains. Second, the promoter and transgene are flanked by LoxP (5’) and Lox2272 (3’) sequences to enable future users to swap in a different transgene at a known integration site via Cre-mediated recombination. Third, the complete insert, consisting of lox sequences, the LvP promoter, and the ENG3K transgene, is flanked by Minos-associated inverted terminal repeat (ITR) sequences to enable Minos transposase-mediated transgenesis. Overall, this design allowed for visual screenability during the generation of the LvP::ENG3K line and inclusion of genetic elements useful for forward-looking applications using transgenic sea urchins.

**Figure 2.**
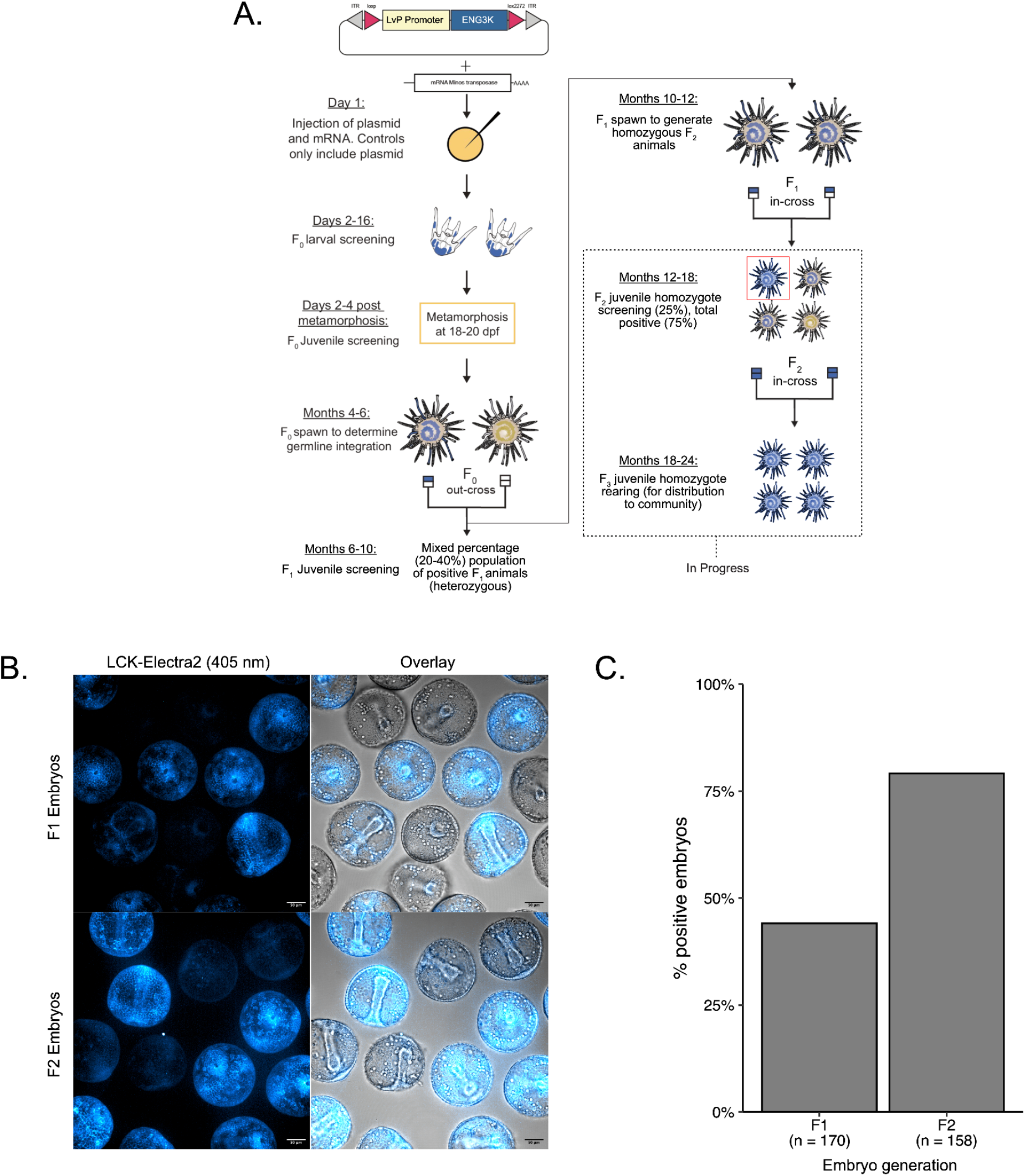
Establishing a sea urchin line for efficiently generating endogenously tagged lines via CRISPR/Cas9-mediated knock-in. **A)** Breeding scheme for the generation of the ENG3K transgenic line. **B)** Confocal images of Electra2 fluorescence in positive F1 generation ENG3K animals and positive F2 generation ENG3K animals **C)** Bar plot showing the proportion of positive individuals in F1 and F2 ENG3K batches.

We established a breeding scheme for achieving the generation of F1, F2 and ultimately F3 homozygous batches of the LvP::ENG3K line (Figure 2A). To establish the line, we injected 1200 wildtype *Lytechinus pictus* eggs with the pUC19-LvP::ENG3K construct and Minos-transposase mRNA. Of these, 538 individuals successfully metamorphosed. Of the 538 F0 juveniles, we found 135 animals which were positive for somatic integrations, and of these positive animals, we qualitatively selected 32 animals which had the most area coverage of fluorescent signal across the test for, and identified two females and two males with germline integration. From these, we selected one male (male #7) with germline integration, that produced non variegated offspring. We then generated F1 animals by outcrossing male #7 with a wild-type female. Following the expected Mendelian proportions (50%), 75/170 (44.1%) of the batches of F1 embryos possessed the transgene (Figure 2B, 2C). Positive F1 adults were screened, sexed, and mated to produce inbred F2 LvP::ENG3K animals. Again following expected ratios (75%), 125/158 embryos (79.1%) of F2 embryos possessed the transgene (Figure 2B, 2C).

Next, we sought to test the split-FP approach across multiple gene targets in order to evaluate its suitability to produce diverse knock-in lines without any subcellular localization constraints. These additional targets included the biomineralization-associated 19kDa protein (*P19*), polyketide-synthesizing enzyme polyketide synthase 1 (*PKS1*), and *polyubiquitin-C*, each representing distinct expression domains during early sea urchin development. A cocktail of Cas9 protein, sgRNA, and ssODN were microinjected into F2 LvP::ENG3K embryos (Figure 3A) and screened at 24 hpf (Figure 3B). The insertion of the small mNG fragment into these endogenous loci resulted in clear fluorescence in the expected cell populations, including pigment cells (*PKS1*), gut/endoderm cells (*FoxA*), and primary mesenchyme cells (*P19*), as observed in both vegetal and lateral views of live embryos.

**Figure 3.**
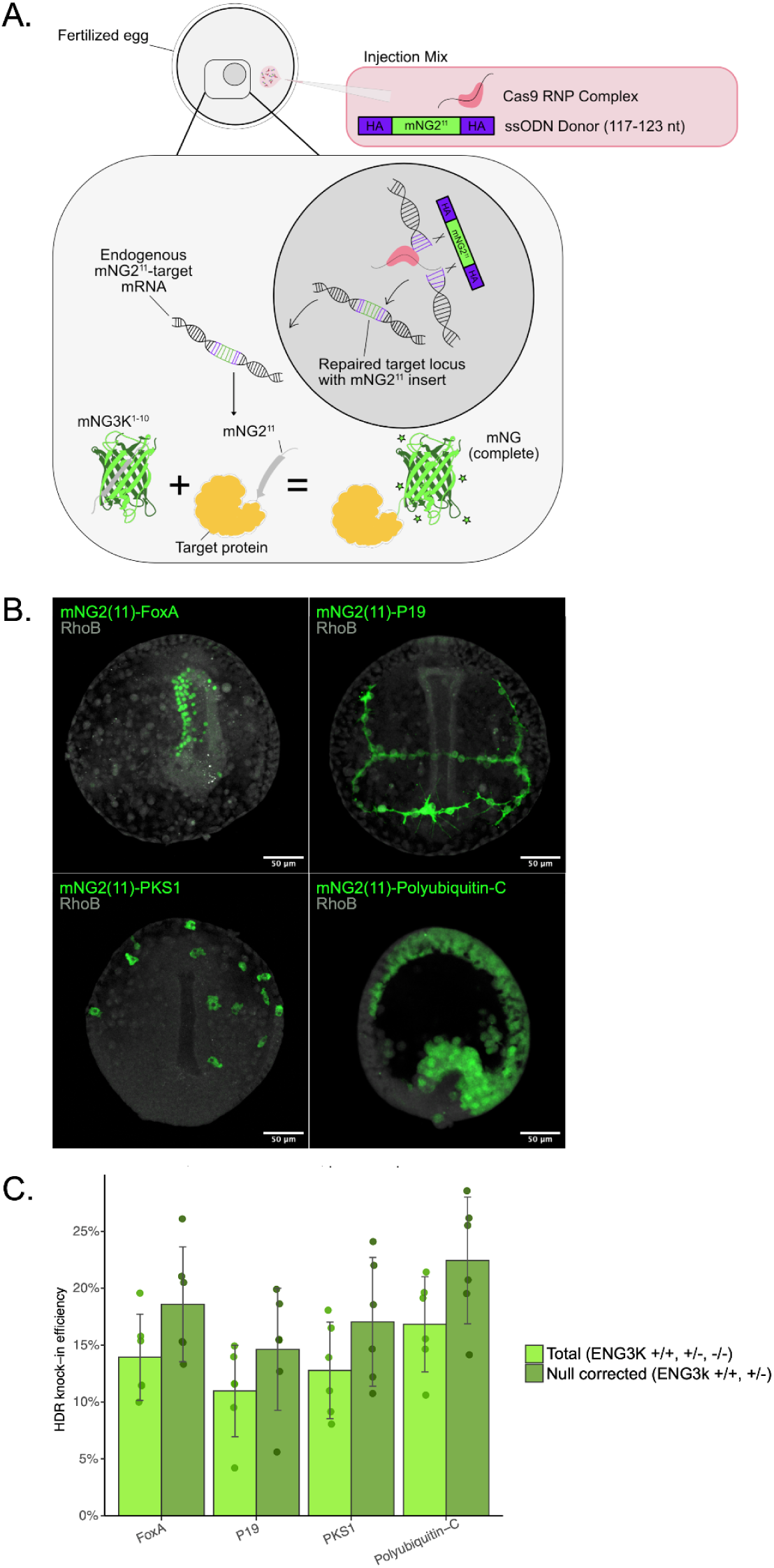
HDR knock-in efficiency of mNG2^11^ at FoxA, P19, PKS1, and Polyubiquitin-C loci. **A)** A schematic illustrating the techniques, reagents, and mechanism of genomic integration using the ENG3K transgenic line for the generation of CRISPR knock-in transgenic lines. **B)** Confocal images of positive individual embryos with integration of the mNG2^11^ sequence at the target locus. **C)** Bar plot with 85% confidence intervals showing the mean efficiency across six technical replicates of each CRISPR knock-in target. Individual overlaid points represent biological replicates.

Mixes were also injected into F1 LvP::ENG3K eggs, fertilized with F1 LvP::ENG3K sperm. Samples were quantified and imaged using confocal microscopy. The mean efficiencies, across six technical replicates and using the same mate pair for each replicate were (Figure 3C): *FoxA* 13.9%, *P19* 10.9%, *PKS1* 12.7%, and *polyubiquitin-C* 16.8%. After accounting for the fact that 25% of animals in the F2 cross are null for the transgene, the adjusted mean efficiencies (adjusted by removal of 25% of the total population) for each target (Figure 3C) were calculated to be, *FoxA* 18.6%, *P19* 14.6%, *PKS1* 17.1%, and *polyubiquitin-C* 22.5%. These results validate the use of a transgenic LvP::ENG3K line as an “acceptor” for CRISPR knock in. Data for each replicate for each target performed for these comparisons are available in Supplementary Table S2.

## Discussion

Echinoderms such as sea urchins have played a major role in developmental biology due to their optical transparency, synchronous development, and accessible embryogenesis. A particularly valuable feature of this organism is their high fecundity, allowing researchers to obtain 0.5-5 x 10^6^ of eggs from a single female, which can easily be fertilized externally and used to generate a virtually unlimited supply of embryos. These features make genetically modified and transgenic sea urchins a promising model for highly accessible and reproducible high-throughput, high-content cell biology (Jackson et al., 2024; Tjeerdema et al., 2023; Vyas et al., 2022).

The results of this study pave the way for CRISPR Cas9-mediated knock-in to become an important complement to existing transgenic approaches in sea urchins. While semi-random integration methods, such as transposon-based systems, enable the generation of reporter lines using exogenous promoters, they are inherently limited by positional effects and the challenge of recapitulating endogenous gene regulation. In contrast, CRISPR-based knock-in approaches provide precise, locus-specific integration that preserves native regulatory context, enabling accurate visualization of protein expression and function at endogenous levels.

The split-FP system described here bridges these two approaches by combining the scalability and stability of a transgenic founder line expressing the large fluorescent fragment via the transposon approach with the precision of CRISPR-mediated insertion of minimal tags at endogenous loci. As a result, these technologies are highly complementary. Transgenic lines can provide a reusable genetic background, while CRISPR knock-in enables flexible, gene-by-gene labeling within that background. This combined framework establishes a versatile and extensible platform for studying protein dynamics in sea urchins and expands the experimental toolkit available for marine developmental biology.

Despite the advantage of the split fluorescent protein approach for improving knock-in efficiency,CRISPR is inherently gene-dependent and subject to several important limitations. Knock-in efficiency can vary substantially across genomic loci due to differences in guide RNA accessibility (Chung et al., 2020), local chromatin environment (Janssen et al., 2019), and the distance between the Cas9 cut site and the desired insertion site (Schubert et al., 2021), all of which influence homology-directed repair (HDR) outcomes (Paix et al., 2017). Even with small donor templates, HDR remains in competition with non-homologous end joining (Yang et al., 2020) and is restricted to specific cell cycle phases (Lin et al., 2014), contributing to variability in editing success across targets and cell types. In the context of split-FP tagging, additional variability arises from incomplete fluorophore reconstitution, which can lead to reduced signal intensity relative to full-length fluorescent protein fusions (Feng et al., 2017; Zhou et al., 2020). Furthermore, tagging efficiency and functional tolerance are highly dependent on the target protein, as insertion at the N- or C-terminus may disrupt protein folding, localization, or activity in a gene-specific manner. Together, these factors highlight that, while split-FP strategies improve scalability, successful implementation still requires empirical optimization on a gene-by-gene basis and careful validation of both expression and protein function.

The broader value of the LvP::ENG3K line lies in the diversity of transgenic resources and experimental approaches it makes accessible. For studies of development and gene regulatory networks, endogenous knock-in reporters provide a means of defining cell identity (Cameron et al., 1987) that extends beyond the snapshots revealed by situ hybridization or transcriptomics, enabling high-fidelity lineage tracing (Tsuyuzaki et al., 2026) as cells progress into later stages of embryogenesis and juvenile development (Tate et al., 2024).

Similarly, this approach is well suited for answering fundamental questions in cell biology of development, where labeling of native proteins can resolve the role of the protein in rapid developmental events such as the cortical reaction, remodeling of the cell surface, and the activity of the machinery involved in cell division and differentiation.

While this approach obviates the need for identification of a specific promoter for each target transgene, it will still be necessary to identify additional promoters to drive expression of the ENG3K transgene earlier than our existing promoter, which expresses at 8-9 hpf (Jackson et al., 2024). The identification of these “egg promoters” is currently underway, and once a suitable candidate has been identified, a line with an egg-promoter-driven split-FP acceptor line will be generated. Additional lines which extend the use of the LvP::ENG3K line are currently being explored. These include lines which express split-FPs of other colors such as split-wrmScarlet (red; (Goudeau et al., 2021)) and split-Capri (cyan; (Tamura et al., 2021)), which when crossed, will enable multiplexed endogenous protein labeling. Lines which express these split-FPs, driven by cell-type specific promoters, for exploring the effects of broadly expressed housekeeping genes in specific cell lineages are being considered as well.

Perhaps most importantly, given the fecundity, optical transparency, and synchronous development of sea urchin embryos, this approach will further extend the reach of high-content imaging experiments using the urchin (Lee et al., 2025). By prioritizing the generation of well-characterized knock-in founder lines, along with the development of community accessible protocols for using such lines, the field could transition from low throughput biology utilizing wild animals, to reproducible, high-throughput imaging of labeled lab strains. Collectively these transgenic sea urchin lines could establish a new framework for studying cell and molecular biology at scale, leveraging the unique advantages of this animal model for contemporary high-throughput biology.

## Materials and methods

### Animals

Wildtype *Lytechinus pictus* individuals used in this study were obtained from La Jolla, CA and maintained as described previously (Jackson et al 2024).

### Molecular cloning

The construct for generating the LvP-ENG3K line (pUC19-LvP::ENG3K; Addgene #259314) was synthesized via Gibson assembly (NEBuilder® HiFi Assembly Kit, New England Biolabs, Ipswich, MA) of fragments obtained from the following plasmids: Minos ITR and LoxP/Lox2272-containing pUC19 backbone (Minos ITR loxp MCS lox2272; Addgene #218983; (Jackson et al., 2024)), *L. variegatus* polyubiquitin-C promoter (LvPolyUb; Addgene #218982; (Jackson et al., 2024)), and a construct containing the ENG3K transgene synthesized by Twist Bioscience (San Francisco, CA).

### Transient expression and functional validation of the split FP system

Injection mixes were prepared at room temperature in the following manner. First, Cas9 endonuclease (1.5 μL, 62 μM, 15.5 μM final concentration) was complexed with sgRNA (2.0 μL, 50 μM, 16.7 μM final concentration) in a microtube for 15 minutes. Second, ssODN (1.5 uL, 10 μM, 2.5 μM final concentration), mNG3K mRNA (0.5 μL, 600 ng/μL, 50 ng/μL final concentration), and Rhodamine B dye (0.5 uL, 10 mg/mL, 0.8 mg/mL final concentration) were added to the mix, bringing the final volume to 6 μL.

### Generation of the Lv-polyubiquitin-C::LCK-Electra2_PT2A_mNG3K^1-10^ transgenic line (ENG3K)

The ENG3K line was generated using Minos transposon-mediated insertion of *LvP::ENG3K*. This was achieved by coinjecting wildtype *L. pictus* eggs (fertilized with wildtype sperm) with a plasmid construct pUC19-LvP::ENG3K (30 ng/μL) and mRNA encoding the Minos transposase (50 ng/μL). Minos transposase mRNA was synthesized by in vitro transcription using HiScribe® T7 mRNA Kit with CleanCap® Reagent AG (New England Biolabs, Ipswich, MA).

Injected F0 generation animals were raised to sexual maturity, and of these animals, one germline integrated male was used to produce outcrossed F1 generation animals. F1 generation animals were raised to sexual maturity and subsequently incrossed to produce inbred F2 ENG3K progeny.

### Generation of CRISPR-Cas9 mNG2^11^ knock-in lines

All reagents (Cas9 endonuclease, sgRNA, and ssODN) used for the generation of CRISPR transgenic animals were purchased from Integrated DNA Technologies (IDT; Coralville, IA), and all nucleic acid sequences used for this study are available in Supplementary Table S3. sgRNAs for targets were selected based on the proximity of the Cas9 cut site to the intended insertion site such that this distance is minimized. However, the majority of the available sgRNA options did not allow for perfect cut-insertion scenarios, where the cut site was either immediately after the start codon or immediately before the stop codon. Thus, various strategies (outlined in Supplementary Data S1) for ssODNs were implemented to achieve perfect, scarless integration. The overarching objective of each strategy was to manipulate ssODN design such that the homology arms of ssODNs perfectly match the sequences flanking the cut site.

Injection mixes were prepared at room temperature in the following manner. First, Cas9 endonuclease (1.5 μL, 62 μM, 15.5 μM final concentration) was complexed with sgRNA (2.0 μL, 50 μM, 16.7 μM final concentration) in a microtube for 15 minutes. Second, ssODN (1.5 uL, 10 μM, 2.5 μM final concentration) and Rhodamine B dye (1 μL, 10 mg/mL, 1.6 mg/mL final concentration) were added to the mix, bringing the final volume to 6 μL. The injection mix was placed on ice until ready for injection. While all experiments and replicates for this study were performed with fresh mixtures, we have not noticed any substantial changes in effectivity of older mixes stored at 4°C within the span of up to one month.

### Imaging

All animals injected with CRISPR reagents were screened for integration at 24 HPF. Screening was performed on the Zeiss LSM 700 (Zeiss, Oberkochen, Germany) using a 488 nm laser for exciting complemented *mNG3K*^*1-10*^/*mNG2*^*11*^ and a 555 nm laser for RhodamineB. Images of membrane LCK-Electra2 were taken using the Molecular Devices HT.ai automated confocal microscope (Molecular Devices, San Jose, CA) using a 405 nm laser. All images taken for this study used a 20X Plan Apochromatic objective on either microscope system.

## Supporting information

Supplemental protocol

## Acknowledgements

We gratefully acknowledge the support of Karen Ong and the National Sea Urchin Resource Center in line production and NIH ORIP 1R24OD 037824.

## References

Boel, A., De Saffel, H., Steyaert, W., Callewaert, B., De Paepe, A., Coucke, P. J. and Willaert, A. (2018). CRISPR/Cas9-mediated homology-directed repair by ssODNs in zebrafish induces complex mutational patterns resulting from genomic integration of repair-template fragments. Dis. Model. Mech. 11, dmm035352.

Cabantous, S., Terwilliger, T. C. and Waldo, G. S. (2005). Protein tagging and detection with engineered self-assembling fragments of green fluorescent protein. Nat. Biotechnol. 23, 102–107.

Cameron, R. A., Hough-Evans, B. R., Britten, R. J. and Davidson, E. H. (1987). Lineage and fate of each blastomere of the eight-cell sea urchin embryo. Genes Dev. 1, 75–85.

Castoe, T. A., Stephens, T., Noonan, B. P. and Calestani, C. (2007). A novel group of type I polyketide synthases (PKS) in animals and the complex phylogenomics of PKSs. Gene 392, 47–58.

Chung, C.-H., Allen, A. G., Sullivan, N. T., Atkins, A., Nonnemacher, M. R., Wigdahl, B. and Dampier, W. (2020). Computational analysis concerning the impact of DNA accessibility on CRISPR-Cas9 cleavage efficiency. Mol. Ther. 28, 19–28.

Feng, S., Sekine, S., Pessino, V., Li, H., Leonetti, M. D. and Huang, B. (2017). Improved split fluorescent proteins for endogenous protein labeling. Nat. Commun. 8, 370.

Gaglia, G. and Lahav, G. (2014). Constant rate of p53 tetramerization in response to DNA damage controls the p53 response. Mol. Syst. Biol. 10, 753.

Goudeau, J., Sharp, C. S., Paw, J., Savy, L., Leonetti, M. D., York, A. G., Updike, D. L., Kenyon, C. and Ingaramo, M. (2021). Split-wrmScarlet and split-sfGFP: tools for faster, easier fluorescent labeling of endogenous proteins in Caenorhabditis elegans. Genetics 217,.

Jackson, E. W., Romero, E., Kling, S., Lee, Y., Tjeerdema, E. and Hamdoun, A. (2024). Stable germline transgenesis using the Minos Tc1/mariner element in the sea urchin Lytechinus pictus. Development 151, dev202991.

Janssen, J. M., Chen, X., Liu, J. and Gonçalves, M. A. F. V. (2019). The chromatin structure of CRISPR-Cas9 target DNA controls the balance between mutagenic and homology-directed gene-editing events. Mol. Ther. Nucleic Acids 16, 141–154.

Jinek, M., Chylinski, K., Fonfara, I., Hauer, M., Doudna, J. A. and Charpentier, E. (2012). A programmable dual-RNA-guided DNA endonuclease in adaptive bacterial immunity. Science 337, 816–821.

Kamiyama, R., Banzai, K., Liu, P., Marar, A., Tamura, R., Jiang, F., Fitch, M. A., Xie, J. and Kamiyama, D. (2021). Cell-type-specific, multicolor labeling of endogenous proteins with split fluorescent protein tags in Drosophila. Proc. Natl. Acad. Sci. U. S. A. 118, e2024690118.

Lee, Y., Jenniches, C., Metry, R., Renaudin, G., Kling, S., Tjeerdema, E., Jackson, E. W. and Hamdoun, A. (2025). Automated, high-throughput in situ hybridization of sea urchin (Lytechinus pictus) embryos. Development 152,.

Ligunas, G. D., Paniagua, G., LaBelle, J., Ramos-Martinez, A., Shen, K., Gerlt, E. H., Aguilar, K., Nguyen, A., Materna, S. C. and Woo, S. (2024). Tissue-specific and endogenous protein labeling with split fluorescent proteins. bioRxiv 2024.02.28.581822.

Lin, S., Staahl, B. T., Alla, R. K. and Doudna, J. A. (2014). Enhanced homology-directed human genome engineering by controlled timing of CRISPR/Cas9 delivery. Elife 3, e04766.

Liu, D., Awazu, A., Sakuma, T., Yamamoto, T. and Sakamoto, N. (2019). Establishment of knockout adult sea urchins by using a CRISPR-Cas9 system. Dev. Growth Differ. 61, 378–388.

O’Hagan, D., Kruger, R. E., Gu, B. and Ralston, A. (2021). Efficient generation of endogenous protein reporters for mouse development. Development 148,.

Oulhen, N., Morita, S., Warner, J. F. and Wessel, G. (2023). CRISPR/Cas9 knockin methodology for the sea urchin embryo. Mol. Reprod. Dev. 90, 69–72.

Paix, A., Folkmann, A., Rasoloson, D. and Seydoux, G. (2015). High efficiency, homology-directed genome editing in Caenorhabditis elegans using CRISPR-Cas9 ribonucleoprotein complexes. Genetics 201, 47–54.

Paix, A., Folkmann, A., Goldman, D. H., Kulaga, H., Grzelak, M. J., Rasoloson, D., Paidemarry, S., Green, R., Reed, R. R. and Seydoux, G. (2017). Precision genome editing using synthesis-dependent repair of Cas9-induced DNA breaks. Proc. Natl. Acad. Sci. U. S. A. 114, E10745–E10754.

Papadaki, S., Wang, X., Wang, Y., Zhang, H., Jia, S., Liu, S., Yang, M., Zhang, D., Jia, J.-M., Köster, R. W., et al. (2022). Dual-expression system for blue fluorescent protein optimization. Sci. Rep. 12, 10190.

Schubert, M. S., Thommandru, B., Woodley, J., Turk, R., Yan, S., Kurgan, G., McNeill, M. S. and Rettig, G. R. (2021). Optimized design parameters for CRISPR Cas9 and Cas12a homology-directed repair. Sci. Rep. 11, 19482.

Seleit, A., Aulehla, A. and Paix, A. (2021). Endogenous protein tagging in medaka using a simplified CRISPR/Cas9 knock-in approach. Elife 10, e75050.

Shaner, N. C., Lambert, G. G., Chammas, A., Ni, Y., Cranfill, P. J., Baird, M. A., Sell, B. R., Allen, J. R., Day, R. N., Israelsson, M., et al. (2013). A bright monomeric green fluorescent protein derived from Branchiostoma lanceolatum. Nat. Methods 10, 407–409.

Shyu, Y. J., Liu, H., Deng, X. and Hu, C.-D. (2006). Identification of new fluorescent protein fragments for bimolecular fluorescence complementation analysis under physiological conditions. Biotechniques 40, 61–66.

Tamura, R., Jiang, F., Xie, J. and Kamiyama, D. (2021). Multiplexed labeling of cellular proteins with split fluorescent protein tags. Commun Biol 4, 257.

Tate, H. M., Barone, V., Schrankel, C. S., Hamdoun, A. and Lyons, D. C. (2024). Localization and origins of juvenile skeletogenic cells in the sea urchin Lytechinuspictus. Dev. Biol. 514, 12–27.

Tjeerdema, E., Lee, Y., Metry, R. and Hamdoun, A. (2023). Semi-automated, high-content imaging of drug transporter knockout sea urchin (Lytechinus pictus) embryos. J. Exp. Zool. B Mol. Dev. Evol.

Tsuyuzaki, K., Yaguchi, J., Yamamoto, T., Ikeo, K. and Yaguchi, S. (2026). Single-cell transcriptomic resources for tracing neurogenesis and cell fate specification in sea urchin embryos. Development 153, dev205025.

Vyas, H., Schrankel, C. S., Espinoza, J. A., Mitchell, K. L., Nesbit, K. T., Jackson, E., Chang, N., Lee, Y., Warner, J., Reitzel, A., et al. (2022). Generation of a homozygous mutant drug transporter (ABCB1) knockout line in the sea urchin Lytechinus pictus. Development 149,.

Yaguchi, S., Yaguchi, J., Suzuki, H., Kinjo, S., Kiyomoto, M., Ikeo, K. and Yamamoto, T. (2020). Establishment of homozygous knock-out sea urchins. Curr. Biol. 30, R427–R429.

Yang, H., Ren, S., Yu, S., Pan, H., Li, T., Ge, S., Zhang, J. and Xia, N. (2020). Methods favoring homology-directed repair choice in response to CRISPR/Cas9 induced-double strand breaks. Int. J. Mol. Sci. 21, E6461.

Zhou, S., Feng, S., Brown, D. and Huang, B. (2020). Improved yellow-green split fluorescent proteins for protein labeling and signal amplification. PLoS One 15, e0242592.

