## Supplemental protocol for "Efficient Endogenous Tagging in the Sea Urchin, *Lytechinus pictus*, Using CRISPR/Cas9-mediated Split-Fluorescent Protein Knock-In"

**Purpose of this guide:**
The purpose of this guide is to describe the criteria for design of guide RNAs and homology arms for CRISPR mediated integration of the short fragment of a split fluorescent protein (mNG11) into a sea urchin ubiquitously expressing mNG 1-10. The expected outcome of this approach is that fluorescence will be produced exclusively in cells expressing the short fragment which has been integrated by HDR.

The subtabs of this document outline the different scenarios (SC) in which certain ssODN design strategy choices need to be made based on the CRISPR guide RNA cut site. The overarching objective of each strategy is to manipulate ssODN design such that the homology arms of ssODNs perfectly match the sequences flanking the cut site. The cut sites in such scenarios are termed **"engineered perfect cut sites"**. Cut sites in scenarios which require no additional manipulation and design of the ssODN beyond simply concatenating homology arms and the desired insert sequence are termed **"natural perfect cut sites"**. That said, this guide describes fixed size characteristics of guides for our lab. However, modifications may be employed by others if you wish to modify the characteristics of the start or stop regions, linker type, or the smallFP tag (split-mScarlet, split-EBFP, etc.).

Some reading guides/shortcuts/definitions:

- "Homology arms": Sequences which flank the cut site.
- "Insert": Any and all sequence/bases which are between homology arms.
- "mNG2(11)": SmallFP tag; TELNFKEWQKAFTDMM
- SCs 1-4 pertain to CRISPR HDR Knock-in at the N-terminus of a target.
- SCs 5-7 pertain to CRISPR HDR Knock-in at the C-terminus of a target.
- In **red** in each subtab is the primary criterion for selecting a particular scenario. Summary guide for criteria:
  - SC1: The cut site is **immediately after** the start codon sequence ("ATG").
  - SC2: The cut site is either **-1 or -2** from the "G" in the start codon sequence ("ATG").
  - SC3: The cut site is **-3 or greater** from the "G" in the start codon sequence ("ATG").
  - SC4: The cut site is **+1 or greater** from the "G" in the start codon sequence ("ATG").
  - SC5: The cut site is **immediately before** the stop codon sequence ("TAA", "TAG", "TGA").
  - SC6: The cut site is either **+1 or +2** from the "T" in the stop codon sequence ("TAA", "TAG", "TGA").
  - SC7: The cut site is **-1 or greater** from the "T" in the start codon sequence ("TAA", "TAG", "TGA").

SC1: N-terminal perfect cut and insertion

**This scenario will never add any additional amino acids to the insertion besides the desired insert.**

1. **The guide RNA cuts immediately after the start codon sequence (“ATG”).**
2. The sequence for either of the homology arms perfectly matches the respective 30 bases flanking the cut site.
3. The length of the insertion sequence is 57nt (mNG2(11)-GSG/GGG).
4. The sequence preceding the start codon is unmodified.
5. The sequence subsequent to the start codon is unmodified.
6. The linker sequence comes **after** the sequence for mNG2(11).
7. The total length of the insert sequence should **always be 57nt**. The total length of the ssODN should **always be 117nt**.

<u>SC2: N-terminal ATG cut and insertion</u>

**This scenario will never frameshift the protein or add any undesired amino acid at the 5' end of the insert. This scenario will always add either a glycine or methionine at the 3' end of the insert.**

1. **The cut site is either -1 or -2 from the "G" in the start codon sequence ("ATG").**
   a. A cut at the position -1 of the start codon would be a cut at the hyphen here: "AT-G".
   b. A cut at the position -2 of the start codon would be a cut at the hyphen here: "A-TG".
2. The sequence for either of the homology arms perfectly matches the respective 30 bases flanking the cut site.
3. The 5' end of the insert sequence contains additional bases:
   a. "G" for a cut at the position -1 of the start codon.
   b. "TG" for a cut at the position -2 of the start codon.
4. The 3' end of the insert sequence contains additional bases:
   a. "GG" for a cut at the position -1 of the start codon.
   b. "A" for a cut at the position -2 of the start codon.
5. The sequence preceding the start codon is unmodified.
6. The sequence subsequent to the start codon may be modified such that the codon immediately after the insertion sequence codes for either:
   a. glycine (for a -1 cut scenario) or
   b. methionine (for a -2 cut scenario).
7. For either -1 or -2 cut scenarios, the total length of the insert sequence **should always be 60nt**. The total length of the ssODN **should always be 120nt**.

SC3: N-terminal pre-ATG cut and insertion

**This scenario will never add any undesired amino acid at the 5' end of the insert. This scenario will always add at least two undesired amino acids only at the 3' end of the insert, one of them always being a methionine.**

1. **The cut site is -3 or greater from the "G" in the start codon sequence ("ATG").**
2. The sequence for either of the homology arms perfectly matches the respective 30 bases flanking the cut site.
3. The insertion prefix sequence: The 5' end of the insert sequence contains bases which match the sequence between the cut site and the desired insertion site (which is always immediately after the start codon). For example: for the sequence "NNNNNNNNNNNNNNNNN-NNATGG", the cut site is -5 from the desired insertion site. The sequence of interest here is "NNATG", and this sequence will be at the 5' end of the insertion sequence. The insertion prefix sequence for any target will always contain "ATG".
4. The insertion suffix sequence: The 3' end of the insert sequence contains additional bases which bring the CDS back into frame. There are several sub-scenarios to consider based on the cut site:
    a. The cut site is -4 from the "G" in "ATG". This means that the 3' end of the insert sequence will need to have two additional bases to bring the CDS back in-frame. In this sub-scenario, by always making those two additional bases "GG", the codon immediately following the insert sequence will always code for a glycine and will simply extend the linker sequence by one amino acid.
    b. The cut site is -5 from the "G" in "ATG". This means that the 3' end of the insert sequence will need to have one additional base to bring the CDS back in-frame. In this sub-scenario, your choice of the single additional base should code for the following amino acids in the following prioritization order:
        i. Glycine
        ii. Serine
        iii. Neutral amino acid
    c. The cut site is -6 from the "G" in "ATG". There are no other bases to add as the sequence will be in-frame. However, a non-endogenous and undesired amino acid will be part of the CDS.
    d. The cut site is greater than -6 from the "G" in "ATG".
        i. Avoid targets which fall in this sub-scenario.
        ii. Experiment with SpRY-Cas9.
        iii. Accept the addition of one or more undesired amino acids in the CDS.
5. For -4, -5, -6 cut scenarios, the total length of the insert sequence **should always be 63nt**. The total length of the ssODN **should always be 123nt**. Consider appropriate alternative strategies for scenarios which have cut sites beyond -6 bases from the desired insertion site.

<u>SC4: N-terminal post-ATG cut and insertion</u>

**This scenario will add undesired amino acids at the 5’ and 3’ end of the insert. SPECIAL CASE: THE FIRST AMINO ACID AFTER THE START CODON CODES FOR A THREONINE.**

1. **The cut site is +1 or greater from the “G” in the start codon sequence (“ATG”).**
2. The sequence for either of the homology arms perfectly matches the respective 30 bases flanking the cut site.
3. The insertion prefix sequence: The 5' end of the insert sequence contains bases which bring the CDS back in-frame. There are several sub-scenarios to consider based on the cut site:
    a. The cut site is +1 from the “G” in “ATG”. This means that the 5’ end of the insert sequence will need to have two additional bases to bring the CDS back in-frame. In this sub-scenario, your choice of these additional bases should code for the following amino acids in the following prioritization order:
        i. Glycine
        ii. Serine
        iii. Neutral amino acid
    b. The cut site is +2 from the “G” in “ATG”. This means that the 5’ end of the insert sequence will need to have one additional base to bring the CDS back in-frame. In this sub-scenario, your choice of these additional bases should code for the following amino acids in the following prioritization order:
        i. Glycine
        ii. Serine
        iii. Neutral amino acid
    c. The cut site is +3 from the “G” in “ATG”. There are no other bases to add as the sequence will be in-frame. However, a non-endogenous and undesired amino acid will be part of the CDS.
    d. The cut site is greater than +3 from the “G” in “ATG”.
        i. Avoid targets which fall in this sub-scenario.
        ii. Experiment with SpRY-Cas9.
        iii. Accept the addition of one or more undesired amino acids in the CDS.
4. The insertion suffix sequence: The 3' end of the insert sequence contains additional bases which bring the CDS back into frame. There are several sub-scenarios to consider based on the cut site:
    a. The cut site is +1 or +2 from the “G” in “ATG”. This means that the 3’ end of the insert sequence will need to have one additional base to bring the CDS back in-frame. These/this base(s) should be the base(s) that code(s) for the original natural amino acid of the protein before any modifications.
    b. The cut site is +3 from the “G” in “ATG”. There are no other bases to add as the sequence will be in-frame. However, a non-endogenous and undesired amino acid will be part of the CDS.
    c. The cut site is greater than +3 from the “G” in “ATG”.
        i. Avoid targets which fall in this sub-scenario.
        ii. Experiment with SpRY-Cas9.

iii. Accept the addition of one or more undesired amino acids in the CDS.

5. For +1, +2, +3 cut scenarios, the total length of the insert sequence **should always be 60nt**. The total length of the ssODN **should always be 120nt**. Consider appropriate alternative strategies for scenarios which have cut sites beyond +3 bases from the desired insertion site.

SC5: C-terminal perfect cut and insertion

**This scenario will never add any additional amino acids to the insertion sequence besides the desired insert.**

1. **The guide RNA cuts immediately before the stop codon sequence (“TAA”, “TAG”, “TGA”).**
2. The sequence for either of the homology arms perfectly matches the respective 30 bases flanking the cut site.
3. The length of the insertion sequence is 57nt (GSG/GGG-mNG2(11)).
4. The sequence preceding the stop codon is unmodified.
5. The sequence subsequent to the stop codon is unmodified.
6. The linker sequence comes **before** the sequence for mNG2(11).
7. The total length of the insert sequence **should always be 57nt**. The total length of the ssODN **should always be 117nt**.

SC6: C-terminal TAA/TGA/TAG cut and insertion

**This scenario will always add an undesired tyrosine (Y) or cysteine (C) at the 5’ end of the insert. This scenario will never add additional amino acids at the 3’ end of the insert.**

1. **The cut site is either +1 or +2 from the “T” in the stop codon sequence (“TAA”, “TAG”, “TGA”).**
    a. A cut at the position +1 of the start codon would be a cut at the hyphen here: "T-AA".
    b. A cut at the position +2 of the start codon would be a cut at the hyphen here: "TA-A".
2. The sequence for either of the homology arms perfectly matches the respective 30 bases flanking the cut site.
3. The 5' end of the insert sequence contains additional bases:
    a. Add "CA" for a cut at the position +1 of the stop codon.
    b. Add “C” or “T” for a cut at the position +2 of the stop codon.
4. The 3' end of the insert sequence contains additional bases:
    a. "T" for a cut at the position +1 of the start codon.
    b. "TA" for a cut at the position -2 of the start codon.
5. The 5’ end of the insert always adds bases which insert an undesired tyrosine (Y) or cysteine (C) into the endogenous protein.
6. The 5’ end of the insert always includes additional bases to complete a stop codon sequence.
7. For either +1 or +2 cut scenarios, the total length of the insert sequence **should always be 60nt**. The total length of the ssODN **should always be 120nt**.

SC7: C-terminal pre-TAA/TAG/TGA cut and insertion

**This scenario will never add any undesired amino acid at the 5’ end of the insert. This scenario will always truncate the endogenous protein by one or greater amino acids at the 3’ end of the insert and will always end with the stop codon in-frame.**

1. **The cut site is -1 or greater from the “T” in the stop codon sequence (“TAA”, “TAG”, “TGA”).**
2. The sequence for either of the homology arms perfectly matches the respective 30 bases flanking the cut site.
3. The insertion prefix sequence:
    a. If the cut site is in-frame, no need to add additional bases at the 5’ end of the insertion.
    b. If the cut is out-of-frame, add either one or two bases which complete the codon for the endogenous amino acid in which the cut site resides.
4. The insertion suffix sequence: The 3' end of the insert sequence should always include **TAA**.
5. For any cut site in this scenario, the total length of the insert sequence **should always be 60nt, 61nt, or 62nt**. The total length of the ssODN should **always be 120nt, 121nt, or 122nt**.

**General checklist for ssODN:**

1. Is the ssODN for the correct gene for the correct animal species?
2. Do the insertion and any modifications keep everything in-frame such that flanking amino acids reflect the sequence of the desired coding sequence?
3. Do the amino acids around the insertion area (-10/+10) have high similarity to amino acid sequences in the same terminus of other species (L. variegatus, S. purpuratus, etc.)
4. Is the insert in the correct orientation given the insertion site (N vs C terminus)?
5. Is the name of the ssODN sequence file correct in that it matches the location of the insert (mNG2(11)-X for N-terminal insert vs X-mNG2(11) for C-terminal insert)?
6. Does the target folder name match the ssODN design strategy used for the target?
7. Does the length of the ssODN for this target match the length it should be as stated for each strategy?
8. On the IDT ordering sheet, is the ssODN sequence named with the matching ssODN sequence name in Geneious/your internal file name?

**General checklist for gRNA:**

1. Is the correct guide sequence annotated in the associated ssODN file?
2. Is the guide sequence split at +17/+23 in the associated ssODN file?
3. Is the cut site associated with the gRNA annotated correctly in the ssODN sequence file?
4. Is the name of the gRNA sequence the same as the name of the gRNA sequence in the ssODN annotations?
5. On the IDT ordering sheet, is the gRNA sequence 20nt and 5’ -> 3’?
6. On the IDT ordering sheet, is the gRNA sequence named with the matching gRNA sequence name in Geneious/your internal file name?
